# Both DNA binding domains of p53 are required for its ultra-rapid recruitment to sites of UV damage

**DOI:** 10.1101/2020.02.09.938993

**Authors:** YH. Wang, T. Ho, A. Hariharan, HC. Goh, MP. Sheetz, DP. Lane

## Abstract

p53 concentrates at DNA damage sites within two seconds upon UV laser micro-irradiation. Structural analysis shows that this very rapid response requires both the DNA binding and C-terminal domains of p53. This early recruitment response is also PARP-dependent. As mutations within the DNA binding domain of p53, that are commonly associated with cancer also inhibit this rapid binding, we suggest that this is an important initial step for p53 function as a tumor suppressor.

**One Sentence Summary:** p53 is an early responder to DNA damage

## Main text

P53 is seen primarily as an ‘effector’ protein in the DNA damage response (DDR) signaling cascade. It is well known that an increase in p53 levels and nuclear accumulation occurs following DNA damage(*1*). P53 then functions as a critical transcription factor regulating expression of a variety of target genes. P53-dependent activation of cell cycle arrest via p21 and apoptosis via PUMA and Bax constitute the most extensively characterized roles of p53 in response to DNA double-strand breaks (DSBs)(*2*). However, rapid binding of p53 to DNA damage sites is not generally expected.

p53 has been implicated in transcriptionally-independent regulation of different DNA repair pathways like nucleotide-excision repair, base-excision repair and mismatch repair(*3*). P53 is also emerging as an important factor in regulating the balance between homologous repair (HR) and non-homologous end-joining repair (NHEJ)(*4, 5*). Since choice of repair is strongly tied to damage sensing, these findings indicate that there may be an unreported role for p53 at the crossroads of initial damage sensing and downstream recruitment of and/or transcriptional regulation of relevant repair factors. This is substantiated by findings that effects of p53 deficiency cannot be explained by systematic perturbation of downstream DDR or repair factors(*6*).

To investigate the potential of p53 to function as a sensor of DNA damage, we micro-irradiated nuclei with a 355nm laser and monitored p53 recruitment. When RPE1 cell nuclei, in which the p53 locus has been endogenously tagged with a neongreen reporter, were irradiated at a single point, p53 was recruited to the specific site of damage within two seconds (Fig. 1A, D). The magnitude of accumulation was dependent on the laser power and hence extent of DNA damage (Fig. S1A, B). Discernable accumulation of endogenous p53 at damage sites was found in 76% of U2OS cells fixed within 5 min following microirradiation (Fig. 1F, G). Line profile across damage sites showed a clear co-localization of p53 and γH2AX staining (Fig. 1H). The staining of p53 was lost upon p53 depletion by RNAi and the staining of γH2AX appeared more disperse (Fig. 1I). We also found that under conditions of overexpression of wildtype p53, EGFP-tagged p53 was similarly recruited rapidly to damage sites (Fig. 1B, E). This was in contrast to the p53 DNA-binding mutant, R248Q, which failed to localize to damage sites (Fig. 1C). Hoechst 33342 photo-conversion was detectable only with 405 nm laser irradiation, and not with 355 nm laser irradiation, either in addition to or in the absence of 405 nm laser (Fig. S1C-D). We therefore rule out the possibility of p53 rapid accumulation as a result of photo-conversion artifacts.

**Fig. 1.**
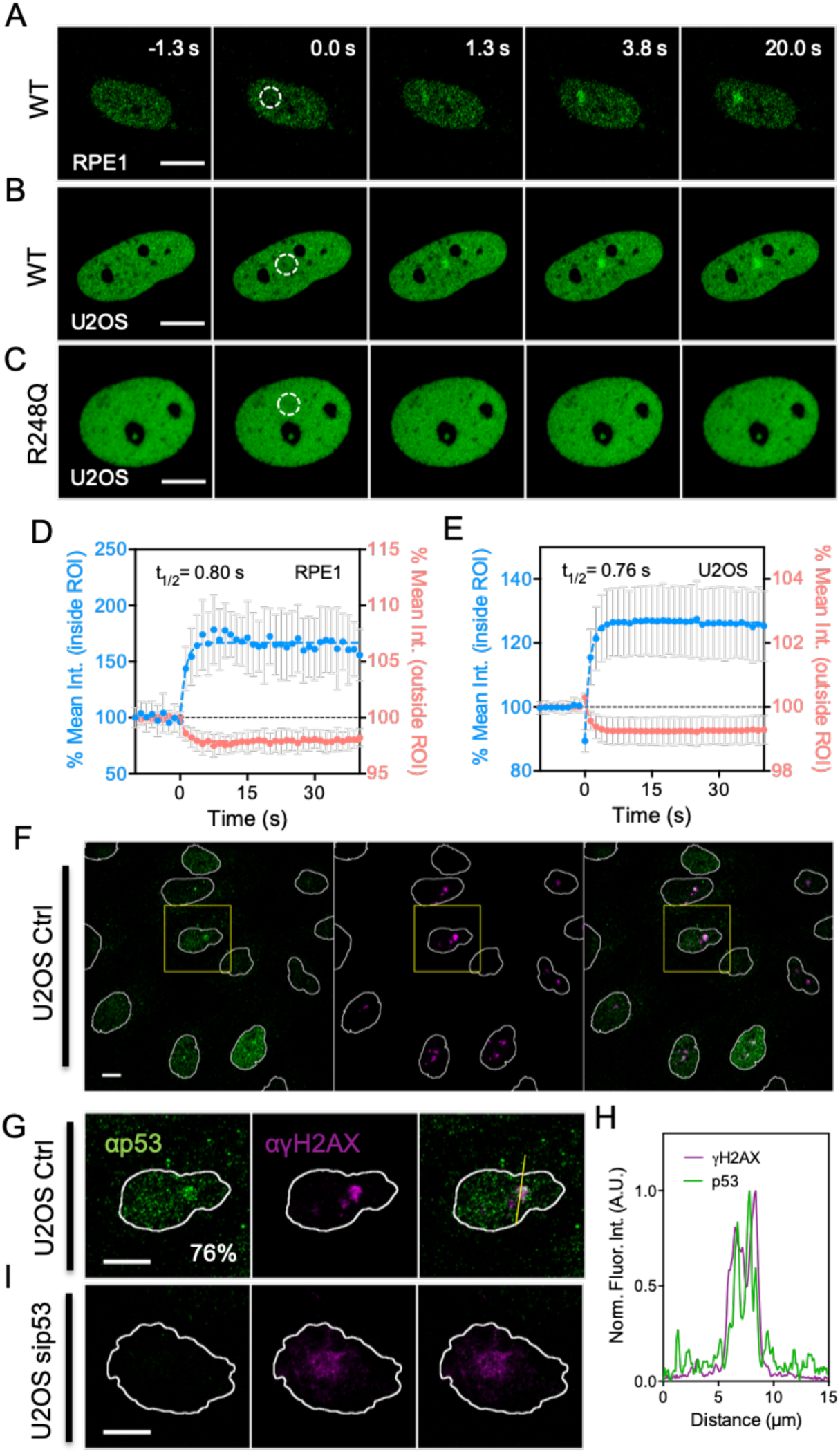
Rapid accumulation of p53 at sites of DNA damage. (**A**) Representative fluorescent micrographs showing the rapid recruitment of endogenous p53 with a CRISPR-KI neongreen tag in RPE1 cells. (**B**) Representative fluorescent micrographs showing rapid accumulation of wildtype (WT) p53-EGFP fusion protein following overexpression in U2OS cells. White dotted circle indicates irradiated area. The diameter of ROIs for quantification is set at 2.5 μm if not indicated otherwise, while the full width at half maxima of the UV laser beam at focal plane was approximately 850 nm. Colored dotted line indicates single exponential fitting. (**C**) The rapid accumulation is not detected with the p53 DNA-binding mutant, R248Q. (**D-E**) Quantification of fluorescence intensity within irradiated area or region of interest (ROI) of (**D**) endogenous p53-neongreen in RPE1 cells (mean±s.d., n=19) and (**E**) WT p53-EGFP in U2OS cells (mean±s.d., n=30). Signal intensity measurements outside the ROI indicate that p53 protein is being redistributed rapidly following irradiation. (**F**) Global and (**G**) zoom in view of immunostaining of endogenous p53 and γH2AX in U2OS cells 5 minutes after irradiation. Yellow line indicates the region used for intensity profile analysis. (**H**) Corresponding fluorescence intensity profile demonstrating co-localization of p53 and γH2AX at early time points after laser irradiation. (**I**) The staining of p53 at DNA damage sites is not detectable in U2OS treated with siRNA against p53 and the staining of γH2AX appears more diffusive.

A measure of intranuclear re-distribution of p53 within and outside of irradiation sites revealed a single exponential time constant of around 0.8 seconds (Fig. 1D, E) and could not be separated from the bleach recovery suggesting that rapid p53 recruitment was diffusion-limited. Together, these data place p53 amongst other early sensors of DNA damage.

TP53 is one of the most frequently mutated genes in cancer, with mutants exhibiting diverse functional consequences which include loss- and gain-of-function(*7*). Mutations in the DNA-binding domain (DBD) of p53 constitute ‘hotspots’ and are commonly identified in human cancers (Fig. 2A). We generated various p53 domain-specific mutants (Fig. 2B, Table S1) and tested their recruitment to sites of DNA damage.

**Fig. 2.**
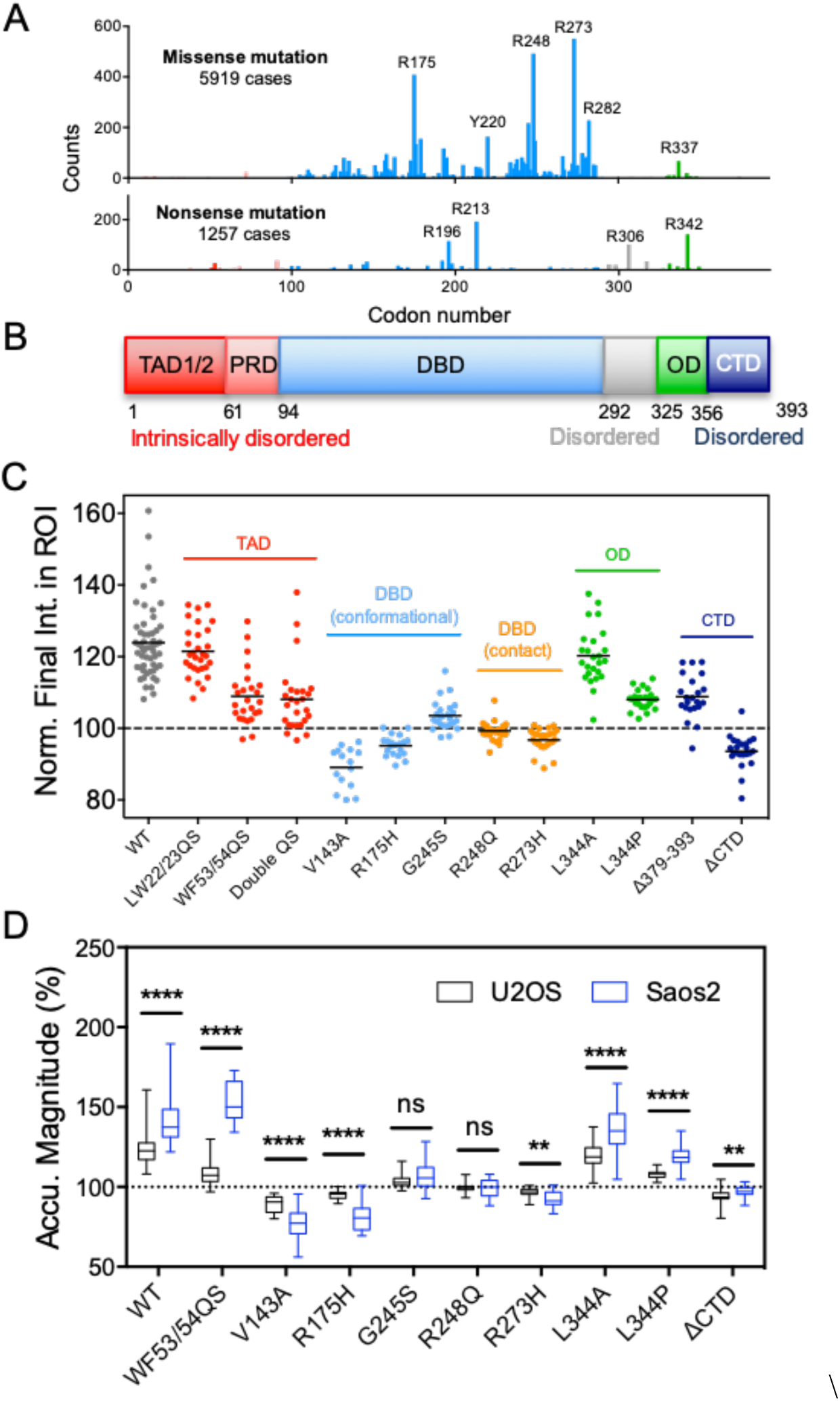
Common p53 mutants in human cancer fail to be rapidly recruited to sites of DNA damage. (**A)** Frequency analysis of missense and nonsense mutations of *TP53* in human cancer based on data available on the COSMIC (Catalogue of Somatic Mutations in Cancer) database. Common ‘hotspot’ areas for mutations are R175, R248 and R273. (**B**) Diagram of human wildtype p53 with various domains indicated. (**C**) Magnitude of accumulation of wildtype (WT) and various mutant p53 as measured by fluorescence intensity within ROI in U2OS cells. (From left to right, n= 54, 18, 20, 18, 30, 21, 25, 26, 24, 19, 15, 22, 20, 25, 31, 30, 27, 25, 25, 25, 23, 21, 22, 23) (**D**) Comparison of the magnitude of accumulation of wildtype (WT) and various mutant p53s in U2OS and Saos2 cells following irradiation.

Strikingly, p53 DBD mutants (conformational (i.e. R175H, V143A, G245S) and DNA-binding contact mutants (i.e. R273H, R248Q)) were defective in recruitment to damage sites and this correlated with the loss of p53 function *in vivo* (Fig. 2C, S2A-B).

We also found that deletion of the CTD (ΔCTD) abolished the early recruitment of p53 to sites of UV damage (Fig. 2C, S2C). Meanwhile, partial loss of the CTD (Δ379-393) partially suppressed p53 recruitment with recruiting speed unaffected. It appeared that both the DBD and CTD were critical in p53 function as a rapid sensor of and responder to DNA damage.

The CTD is also significant as the site for p53 interaction with PARP(*8-10*) and PARylation(*11*). Once bound to p53, PARP can induce PARylation of p53. ΔCTD, which specifically lacked the PAR-interacting domain required for binding to auto-PARylated PARP1, failed to be recruited in both U2OS and Saos2 cells (Fig. 2C, D, S2I). This contrasted with the Δ379-393 mutant, which retains about 20% PARP1-interacting capability and can be PARylated(*11*), and was recruited to a lesser degree.

Examining the N-terminus of p53, we found that mutations in the second, but not the first, TAD disrupted p53 recruitment (Fig. 2C, S2E). In mouse models, the first TAD domain is required for the transcriptional activation of the majority of p53 responsive genes; however the second TAD has been shown to be sufficient for tumor suppression and only loss of both TADs inactivated tumor suppression completely (*12*). In our study, the double TAD (double QS) mutant was also impaired in its recruitment. Thus, mutations that disrupted early recruitment of p53 to DNA damage sites also appeared to strongly correlate with loss of tumor suppression.

P53 functions optimally as a transcription factor as a tetramer. L344A, a mutant that abolished the formation of tetramers but allowed dimers, was recruited to damage sites. Whereas L344P, a mutant which abolished both tetramer and dimer formation, was not recruited (Fig. 2C, S2D).

Our analysis also revealed cell-type dependent differences in the degree of wildtype and mutant p53 recruitment to damage sites. Compared to U2OS cells, which contain wildtype p53, we found a dramatic increase in damage-dependent recruitment of wildtype p53 in Saos2 cells (a p53 null cell line(*13*)) and a slight reduction in three cancer cell lines that expressed either mutant or wildtype p53 (Fig. S2F-G). However, the DBD and ΔCTD mutants consistently failed to be recruited to damage sites in both cell lines (Fig. 2D, S2H).

Given that PARP was a well characterized DNA damage sensor and interacted with p53, we verified rapid recruitment of both these factors in our system. Next, we sought to determine the enzymatic requirements for p53 recruitment. The most profound result was seen with PARP inhibitors which completely blocked the rapid recruitment of endogenous and overexpressed p53 to damage sites in U2OS and Saos2 (Fig. 3B-G, S3A-E). Strikingly, PARP inhibition does not suppress rapid recruitment of PARP (Fig. 3B, C, S3D), suggesting that rapid recruitment of p53 is likely dependent on enzymatic modification by PARP rather than direct interaction with PARP. In stark contrast, ATR inhibition had no effect on p53 recruitment and ATM inhibition showed a more complex phenotype where recruitment was possible, albeit slightly delayed and of lesser magnitude at an initial recruitment (Fig. 3F&G).

**Fig. 3.**
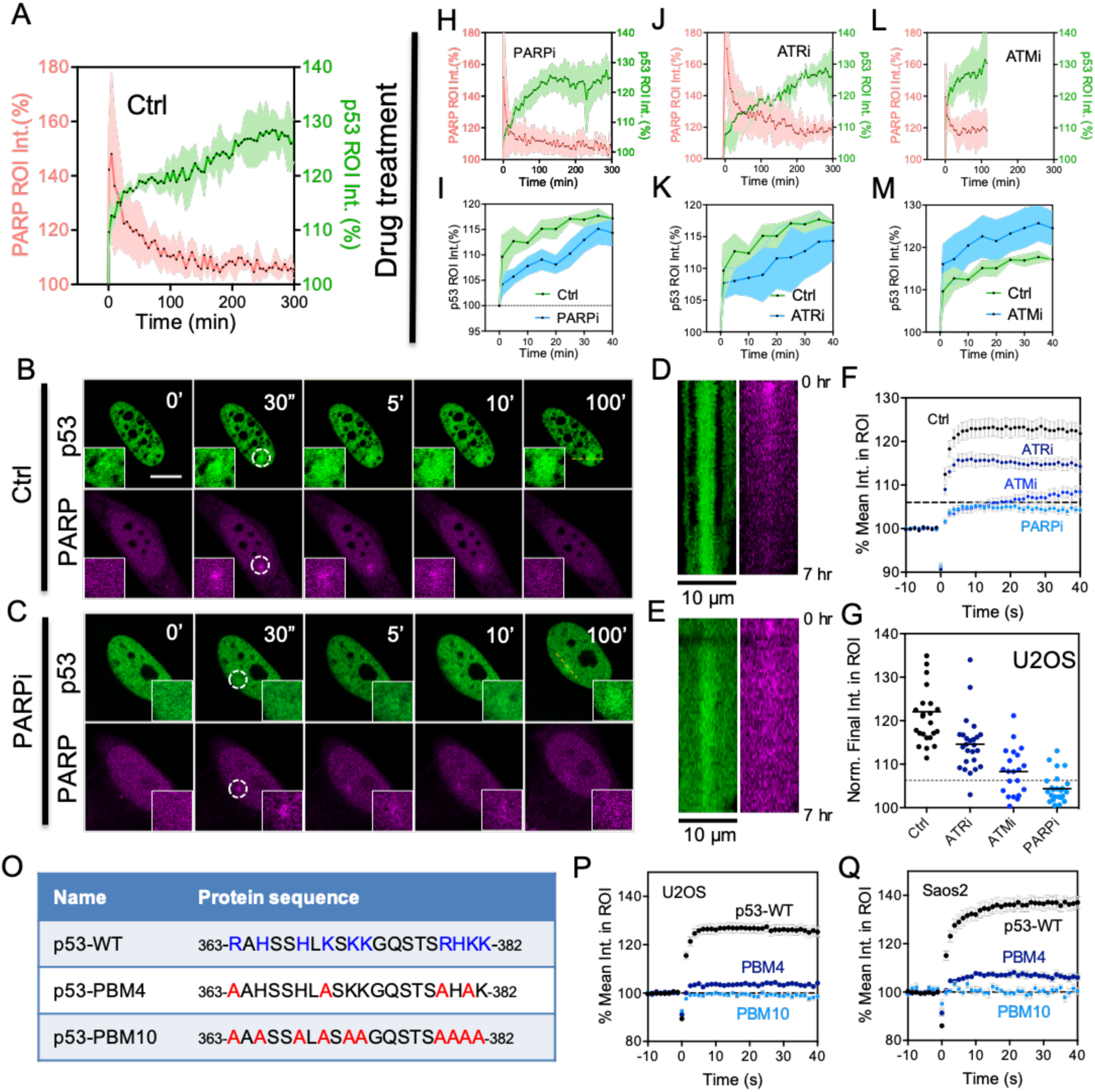
Rapid accumulation of p53 is dependent on PARP activity. (**A**) Quantification of fluorescence intensity within ROI over 300 minutes of wildtype (WT) p53-EGFP and PARP-cb-RFP following irradiation. (**B-C**) Representative fluorescent micrographs showing recruitment of p53-EGFP and PARP-cb-RFP in the first 100 minutes following irradiation, in the absence and presence of PARP inhibition (PARPi), in U2OS cells. (D-E) Kymograph showing different p53 recruitment dynamics following the dotted yellow line showing in panel B&C, respectively. (**F**) Rapid accumulation of WT p53-EGFP in U2OS cells is suppressed or delayed to different extents by various inhibitors (20nM ATRi, 50nM ATMi, 10nM PARPi) as measured within the first 40 seconds following irradiation. (**G**) Magnitude of accumulation of WT p53-EGFP as measured by fluorescence intensity within ROI in the presence of various inhibitors in U2OS cells. (**H, J, L**) Quantification of fluorescence intensity within ROI of WT p53-EGFP and PARP-cb-RFP following irradiation, in absence and presence of various inhibitors, over 5 hours in U2OS cells. (**I, K, M**) Corresponding zoomed in views of F, H, J, focusing on the first 40 minutes of WT p53-EGFP and PARP-cb-RFP recruitment, in absence and presence of various inhibitors, in U2OS cells. (**O**) Protein sequences of PBM4 and PBM10 mutants with disrupted p53-PARP interaction. (**P&Q**) The lack of recruitment of both PBM4 and PBM10 mutants in comparison to the p53-WT in (**P**) U2OS and (**Q**) Saos2 respectively.

**Fig. 4.**
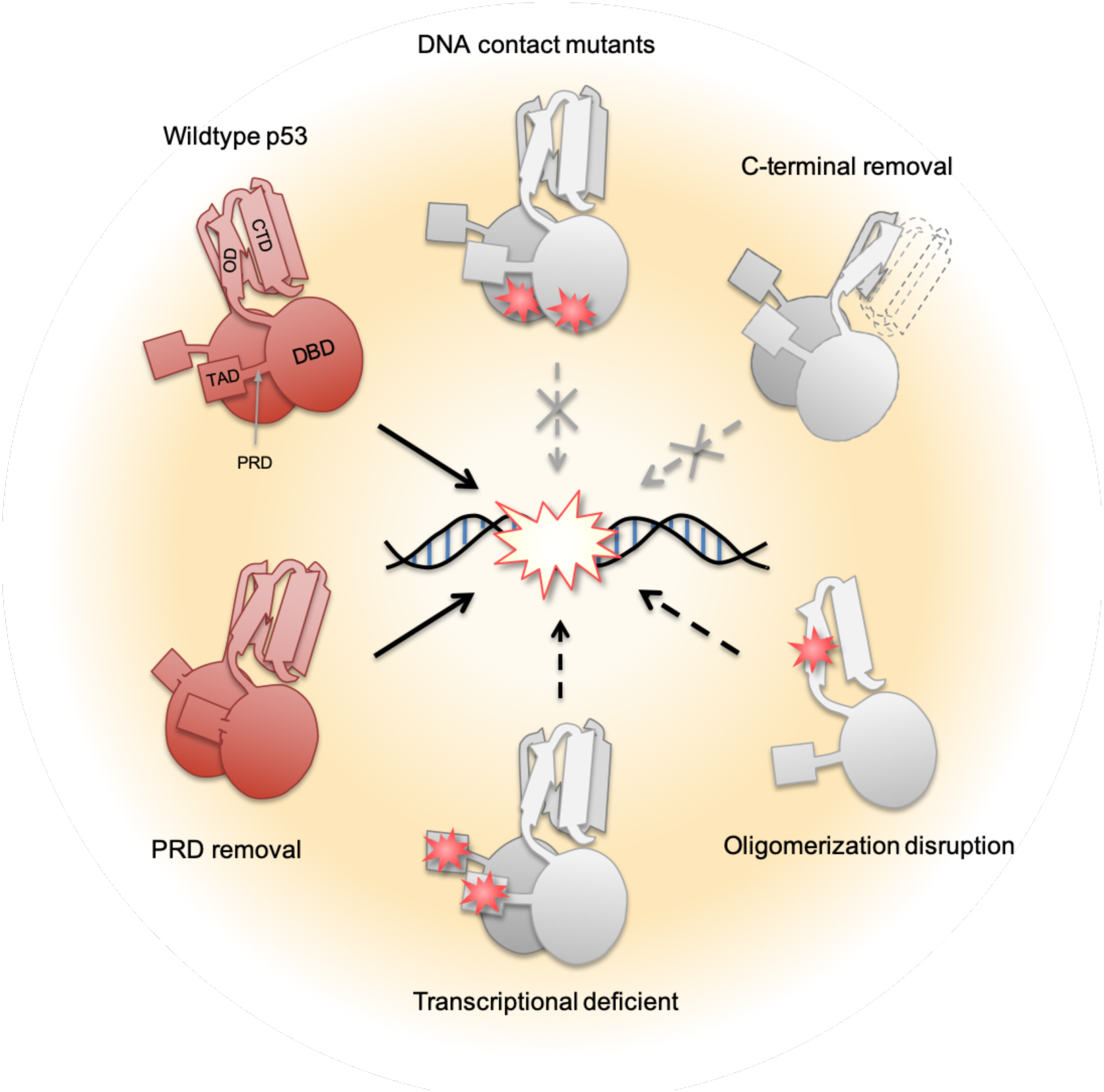
Structural analysis reveals domains required for p53 early recruitment. Solid black arrow represents rapid recruitment comparable to that of WT p53; dotted black arrow represents partial recruitment at a lower magnitude; dotted gray arrow represents failure of recruitment. Red stars represent point mutations in a specific domain.

Based on our above findings and prior studies on the requirements of the CTD for PARylation, we tested PAR-binding-deficient p53 mutants where either four (PBM-4) or ten (PBM-10) critical basic amino acids in the CTD were substituted with alanine(*11*) (Fig. 3O). Their recruitments were also defective (Fig. 3P, Q), suggesting that p53 recruitment was dependent on CTD PARylation.

Here we describe a new property of p53 – its rapid recruitment to sites of DNA damage. The response is remarkably rapid, detected within seconds of UV-irradiation and highly specific. Rapid recruitment of p53 following laser microirradiation mirrors that observed with nuclear phosphoinositides(*14*) which can also complex with p53(*15*). The recruitment kinetics amongst domain-specific p53 mutants also highlights a need for better understanding of the mechanistic implications of such differences on mutant-specific function. Prior work portrays p53 as a late player in the DDR, with protein accumulation, post-translational modification and transcriptional activity occurring hours after damage. Thus, rapid recruitment of p53 indicates that there are early processing events detected by p53 as part of DDR signaling.

## Materials and Methods

### Cell culture

U2OS and Saos2 (human osteosarcoma), SW-13 cells (human primary small cell carcinoma in adrenal gland/cortex), BxPC3 (pancreatic adenocarcinoma) and SK-BR-3 (mammary adenocarcinoma) were obtained from the American Type Culture Collection, ATCC, (catalogue no. HTB-96™, HTB-85™, CCL-105™, CRL-1687™ and HTB-30™ respectively). RPE1 p53-neongreen stable cells were a gift from Karen Oegema. All cell lines except BxPC-3 and SK-BR-3 were maintained in high glucose (4.5 g L^-1^) Dulbecco’s Eagle Medium (DMEM, ThermoFisher Scientific). BxPC-3 cells were maintained in RPMI medium (ThermoFisher Scientific) and SK-BR-3 cells were maintained in McCoy medium (ThermoFisher Scientific), respectively. All media were supplemented with 10% fetal bovine serum (FBS, ThermoFisher Scientific) without antibiotics for cell culture at 37°C in 5% CO2. All cell lines were free from mycoplasma contamination after testing with the MycoAlert™ PLUS mycoplasma detection kit (Lonza) (LT07-701).

### Plasmid constructs and DNA manipulations

Plasmids encoding p53-WT-EGFP was a gift from Tyler Jacks (Addgene #12091). All mutant constructs used in this study were generated from this construct either by a QuickChange II XL kit (Stratagene) or a Q5® Site-directed mutagenesis kit (New England Biolabs) as per the manufacturer’s instructions. PARP1-Chromobody (RFP-tag) plasmid was obtained from Chromotek (#xcr). All constructs were amplified with a Plasmid Midi Kit (Qiagen). All constructs tested in this study, except neongreen-p53, were transiently expressed in selected cell lines using either Neon electroporation transfection system (Invitrogen) or Lipofectamine 2000 (ThermoFisher Scientific).

### Localized DNA damage by laser microirradiation

Localized DNA damage was induced by an UV picosecond laser (355 nm; 1 kHz repetition rate; PowerChip, Teem Photonics) through a 60X objective (NA 1.4, Nikon). Detailed procedure was described elsewhere(*1*). The power of UV laser used in this study ranged 0.2 to 3 nW measured from by the laser scattering light in the optical path, which corresponds to 1 to 15 nW measured at the back aperture of the objective. For convenience, the laser power reported in this study referred to the intensity of the scattering light measured from the optical path, which was set at 0.2 nW unless indicated otherwise.

In short, cells expressing constructs of interest were sensitized with 10 µg mL^-1^ Hoechst 33342 (Sigma-Aldrich) for 10 min. Localized DNA damage was introduced by exposing ROIs within nuclei to a 355nm (UVA) laser for 500 milliseconds at 0.2 nW laser power. The rapid dynamics of p53 recruitment was captured with a single channel live cell imaging with a frame rate at 1.25s per frame with a Galvano scanner, while dual channel videos were taken at a half of the frame rate. The long term observation (up to several hours) were taken at a rate of 5 min per frame. For immunofluorescence analysis, cells were seeded in grid-pattered glass-bottom dishes. Eight to ten cells were selected from each dish with coordinates recorded. UV laser beam were introduced at 20-30 seconds interval including the time needed for alignment so that all cells were ablated and fixed within 5 mins upon laser microirradiation. To investigate the late response of p53, cells were fixed 12 hours after damage. The fixed samples were sequentially stained with primary antibodies against anti-phospho-histone H2A.X Ser139 (Abcam, ab2893) and p53 [clone DO-1] (Cell Signaling Technology #18032); followed by co-staining of secondary antibodies including AlexaFluor488-Goat anti-mouse IgG (A11001, Invitrogen) and AlexaFluor647-donkey anti-rabbit IgG (A31573, Invitrogen). The cells were then imaged again using recorded coordinates with fixed parameters for all samples from the same experiment.

### Image quantification and statistical analysis

To quantify the change in the fluorescent intensity upon laser microirradiation, we first perform image stabilization of all images using FIJI to get rid of obvious translational movement of nuclei, which was usually negligible within a 60 s experiment time window. The UV laser at 1 nW has a full width at half maximum (FWHM) at 0.85 µm measured from the bleaching profile of a sandwiched thin-layer fluorescently labeled agarose gel. Following drift correction using the manual drift correction plugin by ImageJ, a ROI at 2.5 µm in diameter was chosen for analysis, which matches with the size of visible changes in fluorescent intensity due to thermal expansion. The mean fluorescence intensity of the whole nucleus was also quantified and used for bleach correction. To prove that the intensity change within the ROI was due to the redistribution of tagged proteins, we also quantified the mean intensity outside of the ROI but within the nucleus. The data were then processed using a double normalization method following a background subtraction. The data were averaged from experiments with triplicates with about 20-40 cells per condition. Statistical analysis was carried out using GraphPad Prism version 6.04 (GraphPad Software, La Jolla California USA). Two-tailed t-test with Welch’s correction, which does not assume equal standard deviation, was performed throughout this study.

## Acknowledgments

We are grateful to Prof. Yusuke Toyama for his expertise in laser ablation microscopy. We would like to thank various members of the Sheetz and p53 labs for feedback and suggestions.

## Author contributions

YHW, TH, MPS and DPL conceived the ideas, designed the experiments, analyzed the data and wrote the manuscript. YHW and TH conducted the experiments with help from AH and HCG.

## Data and materials availability

All data is available in the main text or the supplementary materials.

## List of Supplementary Materials

Figures S1-S3, Table S1

## References and Notes

1. C. E. Canman et al., Activation of the ATM kinase by ionizing radiation and phosphorylation of p53. Science 281, 1677 (Sep 11, 1998).

2. M. B. Kastan, Commentary on “Participation of p53 Protein in the Cellular Response to DNA Damage”. Cancer research 76, 3663 (Jul 1, 2016).

3. A. Janic et al., DNA repair processes are critical mediators of p53-dependent tumor suppression. Nature medicine 24, 947 (Jul, 2018).

4. S. Moureau, J. Luessing, E. C. Harte, M. Voisin, N. F. Lowndes, A role for the p53 tumour suppressor in regulating the balance between homologous recombination and non-homologous end joining. Open biology 6, (Sep, 2016).

5. T. Rieckmann et al., p53 modulates homologous recombination at I-SceI-induced double-strand breaks through cell-cycle regulation. Oncogene 32, 968 (Feb 21, 2013).

6. V. Menon, L. Povirk, Involvement of p53 in the repair of DNA double strand breaks: multifaceted Roles of p53 in homologous recombination repair (HRR) and non-homologous end joining (NHEJ). Sub-cellular biochemistry 85, 321 (2014).

7. W. A. Freed-Pastor, C. Prives, Mutant p53: one name, many proteins. Genes & development 26, 1268 (Jun 15, 2012).

8. S. R. Kumari, H. Mendoza-Alvarez, R. Alvarez-Gonzalez, Functional interactions of p53 with poly(ADP-ribose) polymerase (PARP) during apoptosis following DNA damage: covalent poly(ADP-ribosyl)ation of p53 by exogenous PARP and noncovalent binding of p53 to the M(r) 85,000 proteolytic fragment. Cancer research 58, 5075 (Nov 15, 1998).

9. J. Wesierska-Gadek, J. Wojciechowski, G. Schmid, Central and carboxy-terminal regions of human p53 protein are essential for interaction and complex formation with PARP-1. Journal of cellular biochemistry 89, 220 (May 15, 2003).

10. H. Vaziri et al., ATM-dependent telomere loss in aging human diploid fibroblasts and DNA damage lead to the post-translational activation of p53 protein involving poly(ADP-ribose) polymerase. The EMBO journal 16, 6018 (Oct 1, 1997).

11. A. Fischbach et al., The C-terminal domain of p53 orchestrates the interplay between non-covalent and covalent poly(ADP-ribosyl)ation of p53 by PARP1. Nucleic acids research 46, 804 (Jan 25, 2018).

12. N. Raj, L. D. Attardi, The Transactivation Domains of the p53 Protein. Cold Spring Harbor perspectives in medicine 7, (Jan 3, 2017).

13. B. Leroy et al., Analysis of TP53 mutation status in human cancer cell lines: a reassessment. Human mutation 35, 756 (Jun, 2014).

14. Y. H. Wang et al., DNA damage causes rapid accumulation of phosphoinositides for ATR signaling. Nature communications 8, 2118 (Dec 14, 2017).

15. S. Choi, M. Chen, V. L. Cryns, R. A. Anderson, A nuclear phosphoinositide kinase complex regulates p53. Nature cell biology 21, 462 (Apr, 2019).

