## Supplementary data for "Both DNA binding domains of p53 are required for its ultra-rapid recruitment to sites of UV damage"

**This PDF file includes:** Figs. S1 to S3**,** Table S1.

**Figure S1**
**Supplementary Figure 1.**

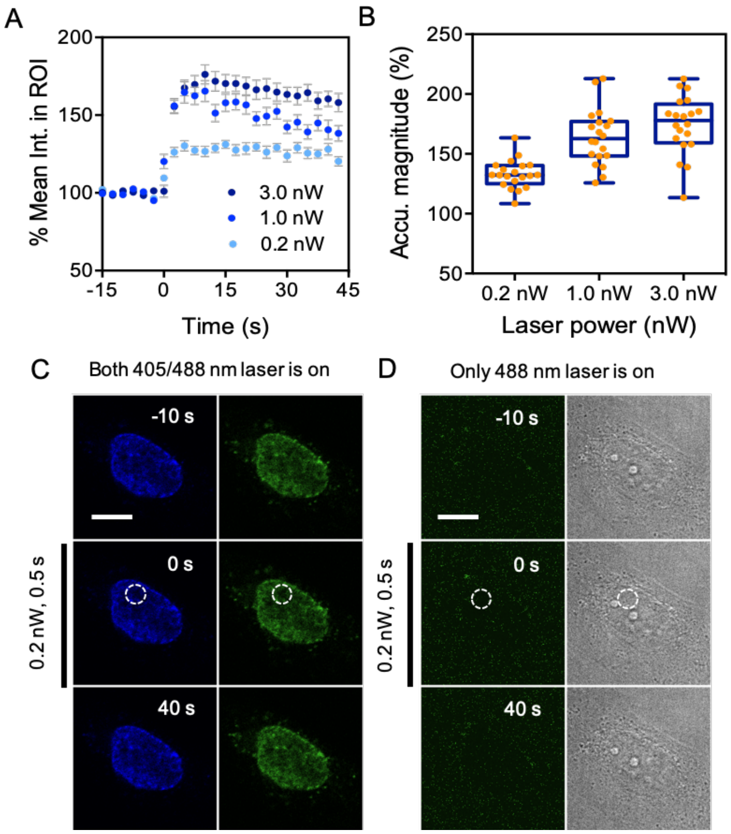


**Fig. S1.** (**A**) The magnitude of p53 early recruitment varies with the 355 nm laser power. (**B**) The maximum magnitude of p53 accumulation within the time course increases with 355 nm power and plateaued at about 1.0 nW. All experiments are carried out at 0.2 nW laser power if not

indicated elsewhere. Microirradiation of the 355 nm laser using Hoechst-sensitized but not transfected U2OS cells doesn't lead to an increased signal in the ROI in the green channel, either (**C**) in the presence or (**D**) absence of the 405 nm irradiation. (**C**) A global Hoechst 33342 photo-conversion is however observed when the cells are being exposed to a 405 nm laser.

**Supplementary Figure 2.**

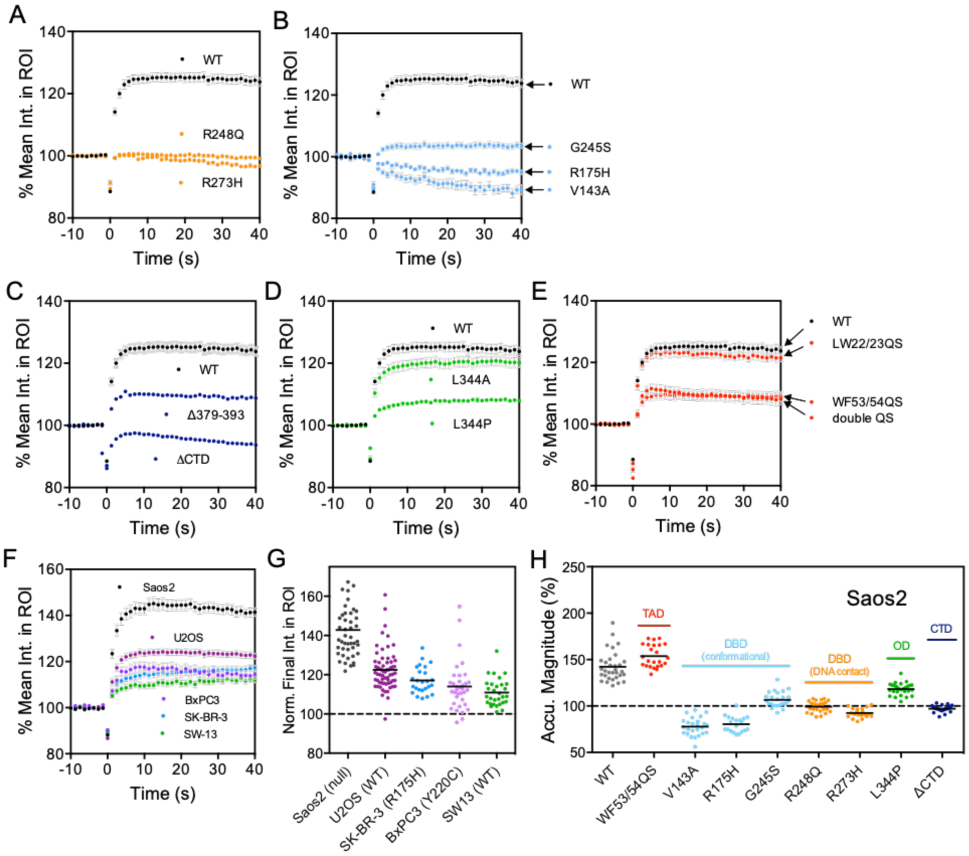


**Fig. S2**. (**A-E**) Time course quantification of fluorescence intensity within ROI of WT and mutant p53-EGFP categorized according to p53 functional domains in U2OS cells. (mean±s.e.m., sample sizes are the same as those indicated in the legend of Fig. 3**D**). (**F**) Time course quantification of fluorescence intensity within ROI of WT p53-EGFP in different cancer cell lines that are null for p53 or contain wildtype or mutant p53. (mean±s.e.m., same sizes for each cell lines from top down are: n= 48, 63, 35, 25, 19, respectively. (**G**) Corresponding quantification of p53 recruitment magnitude at the end of the time course in panel (**F**) (t= 40 s) (**H**) Magnitude of accumulation of wildtype (WT) and mutant p53 as measured by fluorescence intensity within ROI at t= 40 s in Saos2 cells (From left to right, n = 32, 24, 25, 20, 25, 27, 25, 26, 20, 24, respectively) .

**Supplementary Figure 3.**

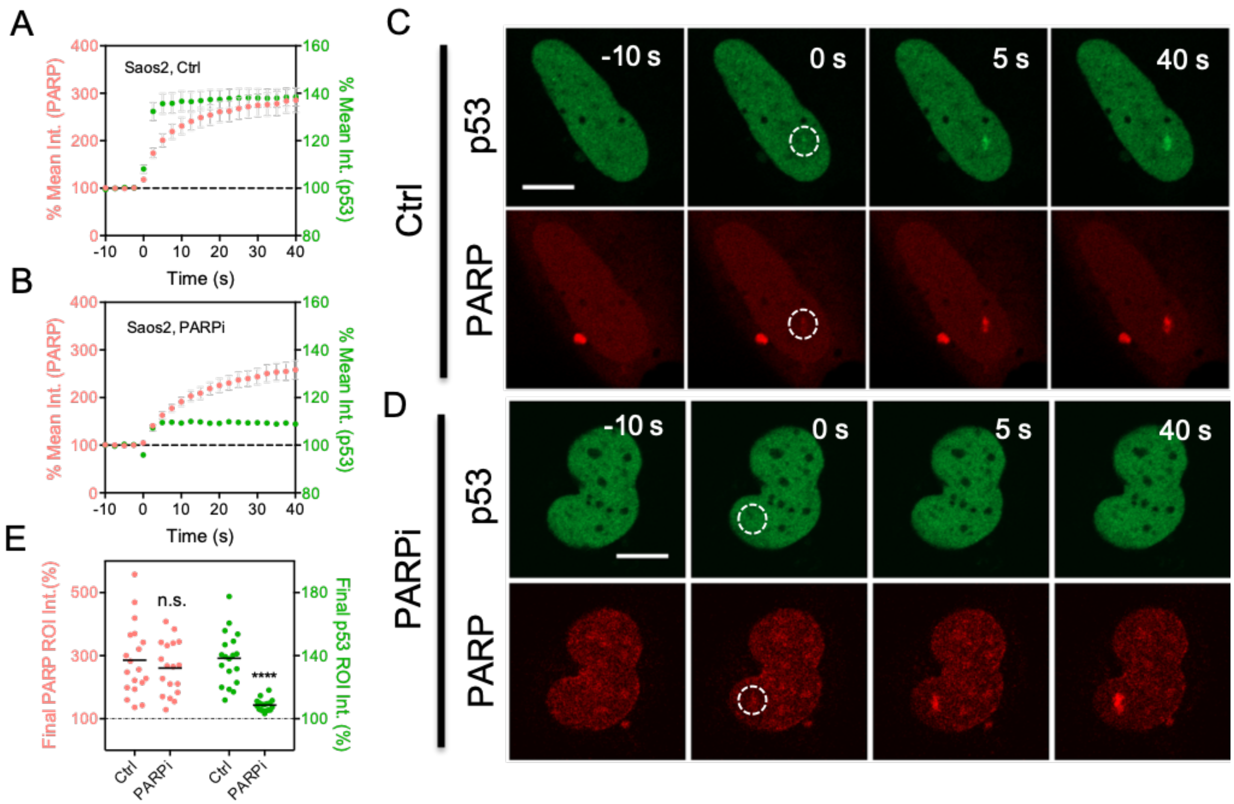


**Fig. S3.** (**A-B**) Quantification of fluorescence intensity within ROI of wildtype (WT) p53-EGFP and PARP-cb-RFP following irradiation over time, in the (**A**) absence and (**B**) presence of PARP inhibition, Veliparib (PARPi), in Saos2 cells. (mean±s.e.m., n= 19 and 18, respectively.) (**C-D**) Representative fluorescent micrographs showing recruitment of p53-EGFP and PARP-cb-RFP in the first 40 seconds following irradiation, in the absence and presence of PARPi. (**E**) Corresponding quantification of p53 and PARP recruitment magnitude at the end of the time course in panel (**A-B**) (t= 40 s).

**Table S1.**

| P53 mutant | Domain targeted for mutation | Remarks | Recruitment |
|---|---|---|---|
| WT | — | Wildtype | Y |
| LW22/23QS | TAD | Disrupt TAD I | Y |
| WF53/54QS | TAD | Disrupt TAD II | N |
| Double QS | TAD | Disrupt both TAD I & II | N |
| V143A | DBD | Conformational mutant | N |
| R175H | DBD | Conformational mutant | N |
| G245S | DBD | Conformational mutant | N |
| R248Q | DBD | DNA contact mutant | N |
| R273H | DBD | DNA contact mutant | N |
| L344A | OD | Dimer mutant | Y |
| L344P | OD | Monomer mutant | N |
| Δ379-393 | CTD | Disrupt p53-PAR interaction | Y |
| ΔCTD | CTD | Abolish p53-PAR interaction | N |

**Table. S1. Summary of domain-specific p53 mutants and recruitment to damage sites. (Y: recruited; N: fail to be recruited; O: partially recruited)**
